# Characterization of enhancer activity in early human neurodevelopment using Massively parallel reporter assay (MPRA) and forebrain organoids

**DOI:** 10.1101/2023.08.14.553170

**Authors:** Davide Capauto, Yifan Wang, Feinan Wu, Scott Norton, Jessica Mariani, Fumitaka Inoue, Gregory E. Crawford, The PsychENCODE Consortium, Nadav Ahituv, Alexej Abyzov, Flora M. Vaccarino

## Abstract

Regulation of gene expression through enhancers is one of the major processes shaping the structure and function of the human brain during development. High-throughput assays have predicted thousands of enhancers involved in neurodevelopment, and confirming their activity through orthogonal functional assays is crucial. Here, we utilized Massively Parallel Reporter Assays (MPRAs) in stem cells and forebrain organoids to evaluate the activity of ∼7,000 gene-linked enhancers previously identified in human fetal tissues and brain organoids. We used a Gaussian mixture model to evaluate the contribution of background noise in the measured activity signal to confirm the activity of ∼35% of the tested enhancers, with most showing temporal-specific activity, suggesting their evolving role in neurodevelopment. The temporal specificity was further supported by the correlation of activity with gene expression. Our findings provide a valuable gene regulatory resource to the scientific community.

**Author summary:** Enhancers are non-coding elements that play a crucial role in the regulation of gene expression during brain development. Despite the availability of various techniques available to identify enhancers, their functional activity is relatively less understood, leaving a gap in our understanding of how enhancer behavior might regulate complex transitions of neurodevelopment. To address this, we utilized forebrain organoids, a 3D model system which closely mimics the complex cellular environment of the developing human brain, and employed Massively Parallel Reporter Assay (MPRA) to validate enhancer activity at various stages of forebrain differentiation, from induced pluripotent stem cells (iPSCs) to neuronal progenitors and cortical neurons. Our study provides a comprehensive catalog of over 2,300 enhancers, showcasing their temporal activity profiles during early neuronal development and offering valuable insights into their likely biological functions. This research advances our understanding of enhancer dynamics in brain development and offers new avenues for further investigations in this field.

## Introduction

It has been over 40 years since the discovery of the first DNA sequence capable of enhancing the transcription of a reporter gene [1]. Since then, many *cis*-regulatory DNA elements known as enhancers have been identified, and their biochemical and functional properties have been extensively investigated. A central role of enhancers is to regulate gene expression by binding transcription factors (TFs) and other regulators able to modulate the transcription of target genes [2]. Enhancers can act independently of the distance and orientation to their target genes [3] and they have specific chromatin features that aid in their genome-wide identification through high-throughput methods. Common techniques used for enhancer identification are DNase I hypersensitivity sequencing (DNase-seq) [4] and the Assay for Transposase-Accessible Chromatin sequencing (ATAC-seq) [5] which exploit the fact that enhancers are free from nucleosomes and are more sensitive to enzymatic treatment. One powerful tool for identifying active enhancers is Chromatin immunoprecipitation-sequencing (ChIP-seq), which utilizes antibodies that recognize specific histone modifications, such as H3K27Ac and H3K4me1, in enhancers flanking nucleosomes [3, 6, 7]. Additionally, active enhancers were found to be actively transcribed into enhancer RNAs (eRNAs) [8, 9], and to form loops with target promoters to exert their regulatory effects on gene expression. In recent years, several high-throughput 3D chromatin conformation techniques, such as Hi-C [10, 11], ChiA-PET [12, 13], and micro-C [14], have been used to identify physical interactions between potential enhancers and promoters, elucidating the spatial organization of the genome.

While these approaches are highly informative and allow identifying the location of potential enhancers, none of these represent the definitive proof of their activity. To address this, reporter assays have been widely used to confirm the activity of predicted enhancers by testing their ability to drive expression of a heterologous reporter gene. Previously, this method was limited to testing one sequence at a time using transgenic technologies [15, 16], but the development of high-throughput massively parallel reporter assays such as STARR-seq [17] and MPRA [18-21] has revolutionized the field, enabling the simultaneous testing of thousands of DNA sequences for enhancer activity in a single experiment.

Here, we used a genomic-integrated MPRA, the Lentiviral-based Massively Parallel Reporter Assay – LentiMPRA; [22] – to evaluate the activity of approximately 7,000 putative gene-linked enhancers. These enhancers were previously identified in human induced pluripotent stem cells (iPSC)-derived forebrain organoids and human fetal cortex combining ChIP-seq and Hi-C data [23] and are potentially involved in early human neurodevelopment.

## Results

### Enhancer selection and lentiMPRA experimental design

To characterize the activity of a set of enhancers putatively involved in early human neurodevelopment, we used lentiMPRA in forebrain organoids to test the activity of a subset of the 96,375 gene-linked enhancers identified by ChIP-seq in our previous study conducted in human fetal cortex and human forebrain organoids [23]. In the original dataset, enhancers were discovered by ChIP-seq, annotated by ChromHMM and linked to putative target genes by proximity and/or external DNA conformation (Hi-C) dataset from stem cells and fetal brains [23]. We generated an MPRA library selecting the most active (e.g., with highest H3K27Ac peak signal) 6,989 enhancers in organoids as measured by ChIP-seq data (**Fig 1A**). Additionally, we included a total of 122 positive control sequences from three different datasets: i) 87 enhancers from human embryonic stem cells (hESCs) validated using the ChIP-STARR-seq approach [24]; ii) 21 MPRA-validated enhancers from hESC-derived neuronal progenitor cells (NPCs) [25]; iii) 35 human brain enhancers from the Vista Enhancer Browser validated by mouse transgenesis [26]. We also included 150 negative controls generated by shuffling the nucleotides of 150 randomly selected candidate enhancer regions. To address the length limitation of oligonucleotide synthesis, we selected a minimal enhancer region of 270 bp by intersecting our enhancers with synonymous information from other datasets, such as p300 ChIP-seq peaks from neuronal cell lines and fetal brain [27, 28], DNA hypersensitivity peaks [29] and CAGE analysis from fetal brain and neuronal cell types [28, 30-32]. The goal was to identify regions with the highest overlap across these datasets which likely corresponds to the core active enhancer region. If the resulting core regions still exceeded the length of 270 bp, we used FIMO [33] to refine them by identifying a subregion with the highest number of known transcription factor binding sites (TFBSs) (see **S1 Fig** and **Methods**). In total, the library included 7,261 sequences synthetized along with 15 bp adapters on either side. The lentiMPRA library was amplified, and a minimal promoter and 15 bp random barcode were placed downstream of each synthesized sequence and cloned into a lentiMPRA vector upstream of the GFP coding sequence (**Fig 1A**).

**Fig 1.**
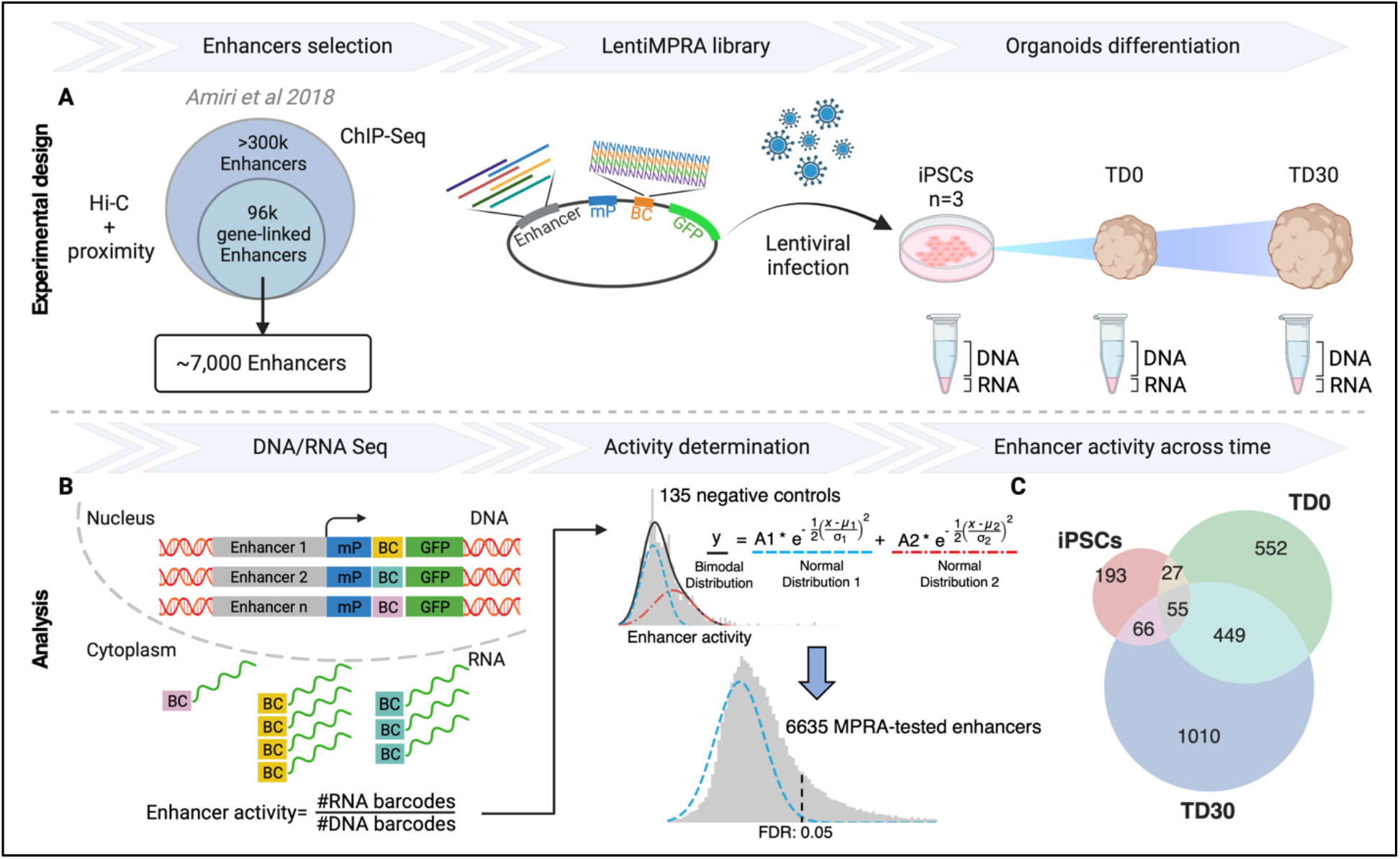
Experimental design and overall lentiMPRA results. **A**. ∼ 7,000 candidate enhancer regions were selected among the 96,000 gene-linked enhancers identified by Amiri et al [23]. The lentiMPRA library was synthesized on a custom array and cloned into a lentiMPRA vector, packaged into lentivirus, and introduced into 3 iPSC lines before organoid differentiation. **B**. At 3 different timepoints, DNA/RNA sequencing was used for estimating the enhancer activity from the ratio of corresponding barcodes. Enhancers were considered active by MPRA if their activity was significantly above the background (blue curve) derived from a gaussian mixture model of the activity of negative controls. **C**. The Venn diagram shows counts of MPRA-active enhancers across time points. BC, barcode; LTR, long terminal repeat; mP, minimal promoter; TD, terminal differentiation.

To investigate whether our candidate enhancers can elicit a time-dependent transcriptional response during early neurodevelopment, we infected three induced pluripotent stem cell (iPSC) lines with our lentiMPRA library and measured enhancer activity at the iPSC stage and at two terminal differentiation (TD) points during forebrain organoid differentiation: an earlier stage (TD0) when organoids were almost exclusively composed of proliferating progenitors and a more mature stage (TD30) when organoids were still harboring progenitors and actively generating layer 5-6 cortical postmitotic neurons (**S2 Fig; Methods**).

### LentiMPRA identifies active enhancers during forebrain organoids differentiation

Using DNA sequencing, we discovered that 95.1% (6,907/7,261) of the tested enhancer sequences in the original library were successfully recovered, with an average of 39.9 unique barcodes associated with each enhancer sequence (**S3 Fig**). Out of the total 7,261 sequences, 94.9% passed stringent quality control (**Methods**). In the MPRA experiment, enhancer activity is measured as the ratio of transcribed barcode reads (obtained by RNA-seq) to integrated genomic barcode reads (evaluated by DNA-seq). To identify active enhancers in the tested set, we first defined the background distribution for the ratio of RNA/DNA barcodes. We reasoned that the presence of a multitude of TFs in each cell, combined with the non-deterministic (i.e., probabilistic) nature of TF binding motifs and the largely unstudied effects of TF cooperativity, could result in enhancer activity even in randomly shuffled negative control sequences. Consequently, the distribution of activity for the negative control sequences was approximated as a sum of two gaussian distributions, one representing the true background signal and one representing an actual signal from potentially active sequences (**Figs 1B and S4; Methods)**. Such a bimodal approximation precisely described the observed experimental distribution with an average signal from likely active sequences being roughly 55% stronger that the average background (**Fig 1B**). Such an approximation also described the signal distribution for positive controls, suggesting that some of the positive controls are not active, albeit enriched for active enhancers as compared to negative controls (**S4 Fig**).

MPRA-active enhancers were defined as those having a signal significantly above the background distribution (**Fig 1B**). Altogether, 34.8% enhancers were active in at least one time point (**S1 Table**). Of the core validated enhancers, 1,755 (74.6%) exhibited activity at a specific time point reflecting a stage-specific epigenomic regulation: 193 were active only in iPSCs, 543 at TD0, and 1,010 at TD30. More than 25% of active enhancers were shared between time points, with a large proportion being active at both TD0 and TD30 (449), indicating that some regulatory elements may play a role in both early and late stages of neurodevelopment **(Fig 1C)**.

The background and the actual signal distribution were wide and overlapped significantly, limiting a clear discrimination between active and inactive enhancers, and reducing the power of the MPRA approach (**Fig 1B**). Based on this overlap and a selected p-value threshold, we estimated that we were only powered to validate about 35% of truly active enhancers. To elaborate on the missing fraction of true enhancers, we clustered all tested enhancers by their activity profile (RNA/DNA ratio) across samples in the MPRA experiment. This clustering approach revealed that almost all enhancers fell into two large clusters, cluster 1 and 2 (**Fig 2A**). Cluster 1 had just a few MPRA-active enhancers, while the cluster 2 encompassed almost all MPRA-active enhancers. We interpreted the data as suggesting that the inactive enhancers in cluster 2 were likely validated, but they did not formally pass the significance test due to the low sensitivity of the MPRA assay. This result would also explain the limited reproducibility of MPRA-active enhancers across samples at TD0 and TD30 (**S5 Fig**).

**Fig 2.**
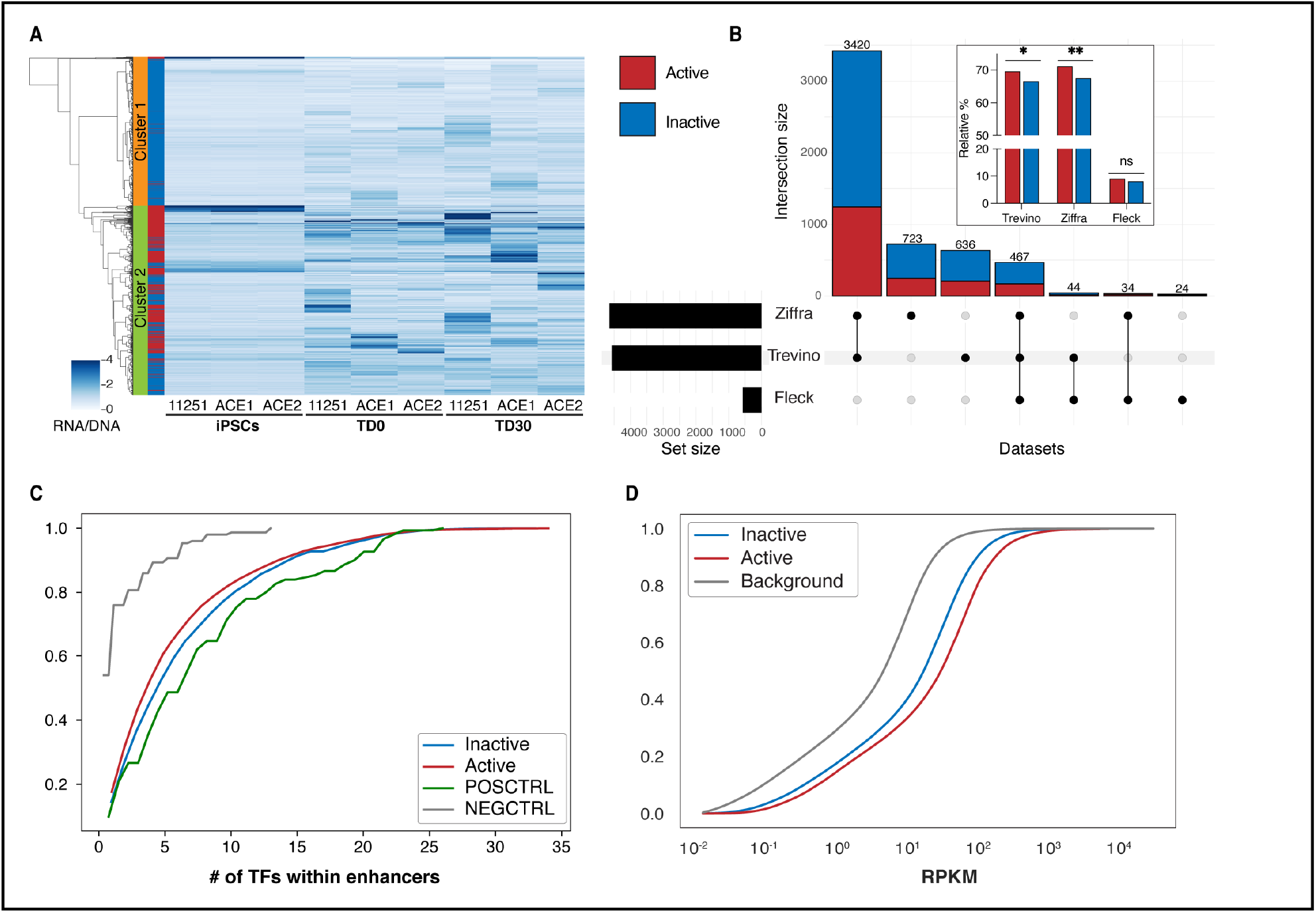
Characteristics of active enhancers. **A**. Heatmap of enhancer activity (RNA/DNA) of all candidate enhancers tested in MPRA experiment. Clustering was performed using Ward variance minimization algorithm. Right bar annotates MPRA-active (red) or inactive (blue) enhancers. **B**. Upset plot showing the number of active (red) and inactive (blue) enhancers overlapping external datasets. In the box, bars represent the fraction of overlapping active and inactive enhancers relative to the total number of active and inactive enhancers. P-values were calculated using Fisher’s exact test (*p-value < 0.05; **p-value < 0.01). **C**. Cumulative plot of number of transcription factor binding sites identified by FIMO in positive controls, negative controls, and tested enhancers. No statistical significantly difference was observed between active and inactive enhancers. **D**. Cumulative plot of the expression (RPKM) of genes predicted to be regulated by tested enhancers in TD0 and TD30 organoids in the endogenous genomic context by Amiri et al. [23]. Genes regulated by MPRA-active enhancers have significantly higher expression than inactive enhancers. The background curve represents the expression of all ∼ 96,375 gene-linked enhancers from Amiri et al [23].

We next intersected MPRA-tested enhancers with external ATAC-seq-derived peaks obtained from bulk and single cell data from fetal brains and forebrain-directed organoids [34, 35] as well as from whole-brain organoids [36]. A large proportion (79.1%) of the tested enhancers were present in these external datasets. Consistent with our expectations, we observed that MPRA-active enhancers were enriched in fetal brains and forebrain-directed organoid datasets from Trevino et al. and Ziffra et al. (**Figs 2B inset, S6** and **S2 Table**).

To further qualify the nature of MPRA-active enhancers, we compared the numbers of TFBSs within active and inactive enhancers. Our analysis revealed that while positive control enhancers had significantly more TFBSs compared to both scrambled negative controls and tested enhancers (**Fig 2C**), we did not observe any significant difference in the number of TFBSs or in the expression of cognate TFs between MPRA-active and -inactive tested enhancers.

We next took advantage of the fact that in the original dataset of Amiri et al. we identified putative target genes for the tested enhancers in their native DNA location, by either proximity in linear DNA and/or 3D DNA conformation datasets [23]. We then compared the MPRA-active and -inactive enhancers with regard to the expression of their linked genes. To increase the statistical power of our analysis, we incorporated bulk RNA-seq data from 88 additional samples from a parallel experiment using genetically distinct iPSC lines (collected at TD0 and TD30, using the same forebrain differentiation protocol) in addition to the samples used in the MPRA experiment. Our analysis revealed that the genes linked to MPRA-active enhancers exhibited significantly higher expression levels than those associated with inactive enhancers (**Fig 2D**) [23]. Genes linked to enhancers in both categories had higher expression than “background”, i.e., all gene-linked enhancers from Amiri et al. [23]. This likely reflects the selection of the most active enhancers for the MPRA experiment. The fact that MPRA-inactive enhancers had higher target gene expression than the “background” expression was consistent with the low sensitivity of MPRA experiment estimated above, implying that a significant fraction of MPRA-inactive enhancers could actually be active in a different experimental setting.

We then correlated the difference in enhancer activity (measured by lentiMPRA) across time points with the difference in expression of the gene(s) linked to the enhancer in the endogenous genomic context. Comparing enhancer activity and gene expression at TD0 versus the iPSC stage, we found that MPRA-active enhancers upregulated in iPSCs or at TD0 were typically correlated (positively or negatively) with the difference in expression of their endogenous linked genes (**Figs 3A, 3B** and **S7**). In contrast, MPRA-inactive enhancers rarely showed a correlation with the expression of their corresponding genes when comparing either TD0 (**Figs 3A and 3B**) or TD30 with iPSCs (**S8 and S9 Figs**). These observations demonstrate that the MPRA-active enhancers, compared to the inactive ones, were those that, in their appropriate genomic context, were linked to highly expressed genes and that differential MPRA activity predicted differential gene expression during differentiation. Given the nature of the MPRA assay, it can be inferred that the activity of these MPRA enhancers is less dependent on the genomic context. Among the enhancers exclusively active in iPSCs, we found that the most active one (**Table S3**) is located upstream of the GTPase-activating protein (SH3 domain)-binding protein 2 (G3BP2) gene, which encodes an RNA binding protein involved in maintaining pluripotency by regulating the transcription factors Oct-4 and Nanog [37] (**Fig 3C**). The G3BP2 gene was also upregulated in iPSCs compared to both TD0 and TD30 stages (**Figs 3A and S8**), consistent with the MPRA results and with its biological role in maintaining pluripotency. The 270 bp tested enhancer region is potentially bound by seven TFs (**S3 Table**), including ZNF263, a TF expressed in pluripotent stem cells [38] and computationally predicted to have the highest affinity for the binding motif (**Fig 3C**).

**Fig 3.**
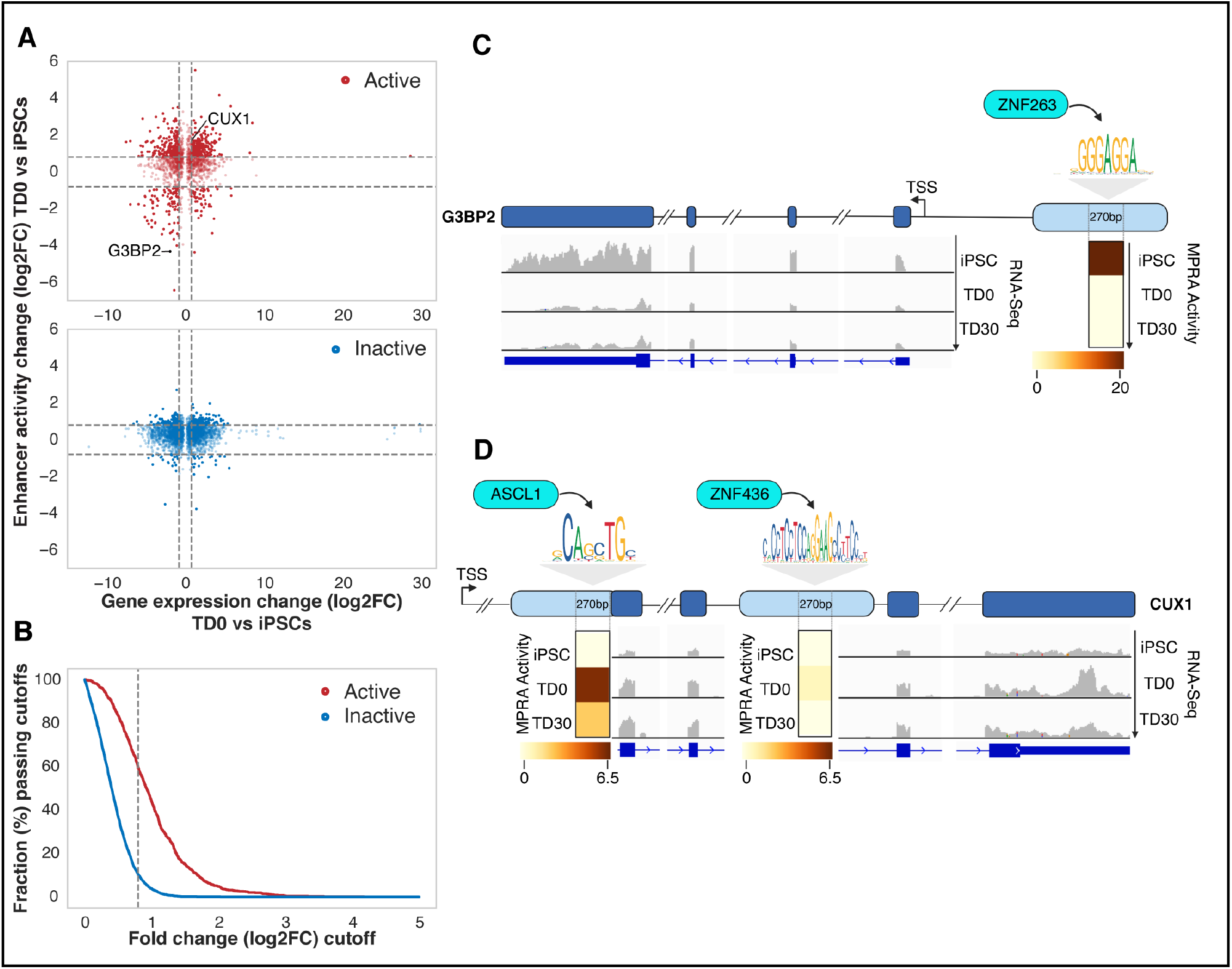
MPRA active enhancers correlate with differences in expression of linked genes across different time points. **A**. Scatter plots of change in MPRA activity for an MPRA-active enhancer (y-axis) and change of expression for the gene(s) linked to the enhancer in the endogenous genomic context. Activity and expression are compared for TD0 and iPSC. Each dot represents a gene-enhancer pair. iPSC or TD0 active enhancers are shown as red circles in the upper panel; inactive enhancers are represented in the bottom panel as blue circles. **B**. More correlated enhancer-gene pairs are observed in MPRA-active enhancers compared to inactive enhancers. The x-axis represents the cutoff for the change in enhancer activity and gene expression. The y-axis represents the fraction of enhancer-gene pairs passing cut off for the change in the enhancer activity and gene expression. The dashed line represents cutoff at log2FC=0.8, which is the same as marked in panel **(A). C**. Example of an iPSC-only active enhancer (MPRA activity shown in the heatmap) and the expression of the linked gene (G3BP2), with ZNF263 as the predicted binding transcription factor. **D**. Example of a TD0 active enhancer (MPRA activity shown in the heatmap) and the expression of the linked gene (CUX1) with ASCL1 as predicted binding transcription factor. For comparison, a mildly active enhancer at TD0 for this gene is identified downstream of the active enhancer. TSS, Transcription Start Site.

One of the most active enhancers, validated both at TD0 and TD30 (**S3 Table**), partially overlapped with an exon located ∼ 400kb downstream of the transcription start Site (TSS) of the Cut Homeobox 1 (CUX1) gene, a TF playing a critical role in upper layer cortical neuron differentiation, dendrite branching, and synapse formation [39, 40]. Among the four TFs predicted to bind this active enhancer (**S3 Table**) there is ASCL1, which has been previously identified to promote CUX1 expression [41]. Additionally, a second intronic enhancer linked to CUX1, bound by ZNF436, showed mild MPRA activity only at TD0 (**Fig 3D**).

## Discussion

Spatial and temporal regulation of gene expression through regulatory regions, specifically enhancers, is essential for shaping the structure and function of the human brain during development. A number of biochemical techniques, such as ChIP-seq, ATAC-seq, DNase-seq have been used for identifying enhancers and characterizing their activity in neurodevelopment. Among those, the MPRAs have been adopted to test an enhancer’s ability to activate a synthetic reporter in a high throughput and context-independent manner, providing an orthogonal readout of enhancer activity. In this study, we used lentiMPRA in forebrain organoids to evaluate, over the course of brain development, the activity of ∼ 7,000 gene-linked enhancers previously identified in human fetal tissues and brain organoids from a combination of histone-mark ChIP-seq and DNA conformational studies. Besides validating 2,352 enhancers, our analysis of the MPRA-active enhancers in relation to upstream binding TFs and downstream targets suggested important implications to the development biology of the human brain.

By analyzing RNA-seq data from forebrain organoids collected at the same time points, we found that while there was no relation with number of potential TFBSs or TFs expression levels, MPRA-active enhancers are associated with highly expressed genes in the endogenous genomic context. Furthermore, differential enhancer activity, as determined by the lentiMPRA assay, tended to be correlated, in a positive or negative fashion, with differential gene expression across timepoints of neural differentiation. This provides an interesting insight into the capability of MPRA to detect enhancers associated with genes dynamically regulated during different stages of neurodevelopment.

On a technical side, our study highlights a few essential limitations of MPRA techniques. Perhaps the major limitation of MPRA is its low sensitivity (poor discrimination of signal from noise), estimated to be about 35% in our experiments, which likely explains limited reproducibility across replicates. For the later differentiation time points this could also be due to lentivirus regulatory element transgene silencing, which was previously reported to be an issue over differentiation of stem cells into other cell types [42]. More data are required to precisely assess the variability of this assay across technical replicates and genetically different iPSC lines. In addition, there may be some false negative results in the MPRA assay, since the assay tests a shorter version of the original enhancer, which may not include all the necessary elements required for transcription initiation. For example, MPRA-tested enhancers may lack crucial co-activators that are provided by DNA loop conformation, or present in the flanking sequences of the tested region. This may explain why, while detecting a significant enrichment of MPRA-active enhancers with external ATAC-seq datasets obtained in fetal brain or forebrain organoids, a considerable percentage of MPRA-inactive enhancers also overlapped with those external datasets.

Similarly to any bulk assay, using MPRA in a heterogenous system such as organoids, which are characterized by cellular diversity, reveals another caveat of the technique. Given the absence of cell-type information coupled to enhancer activity, it is likely that MPRA may be strongly biased towards revealing those enhancers present in the most abundant cell types. Indeed, when intersecting with scATAC-seq data obtained from the same organoids preparations at TD0 and TD30, there was a significant enrichment of MPRA-active enhancers only in the most abundant cell population, radial glia cells (**Table S4**) [43]. Future studies using single-cell MPRA-seq approach [44, 45] in combination with existing single-cell biochemical assays such as scATAC-seq, could open new avenues to gain a comprehensive understanding of the fine regulatory dynamics occurring within each cell type in complex neuronal differentiation systems.

## Materials and Methods

### lentiMPRA library design

Candidate enhancers were identified in Amiri et al 2018 [23] by chromatin segmentation analyses using H3K27Ac, H3K27me3 and H3K4me3 ChIP-seq peaks datasets obtained in cortical organoids and cerebral cortical tissue from postmortem fetal human brains. Enhancers were linked to putative target genes by proximity and/or by association with promoters using fetal brain 3D chromatin conformation data. From an initial dataset of >300,000 enhancers, 96,375 were found to be associated with genes and termed gene-linked enhancers (GLE). From the GLE dataset, approximately 7000 enhancers were selected for MPRA based on top activity, as determined by H3K27Ac peak signal. As ChIP-seq enhancers can encompass hundreds to thousands of bases, for the purpose of oligonucleotide synthesis for the MPRA assay, a core of 270 bp region was identified. To achieve this, our selected enhancers were intersected with enhancers from other datasets to select the region with the highest number of overlaps, and therefore potentially more active. In detail, we used: i) p300 ChIP-seq peaks from human neuronal cell lines [32] and human fetal cortex [46], ii) CAGE analysis from brain tissues [30], and iii) DNase hypersensitivity peaks from neuronal progenitor cells and brain tissue [29]. Finally, if there was no overlap or if the overlapping enhancer region was still too large, we used FIMO [33] to further reduce the size to 270 bp by selecting the region with the highest number of TFBSs, as TF binding is a reliable predictor of enhancer activity. List of tested enhancers is outlined in **Table S1**.

### lentiMPRA library generation

The lentiMPRA plasmid library was constructed as previously described in Gordon et al., 2020 [22]. Briefly, the oligonucleotide pool of the 7,261 enhancers was synthetized by Twist Bioscience and amplified via two rounds of PCR, first to add the minimal promoter and then to add the barcode, using two sets of adaptors primers, 5BC-AG-f01/r01 and 5BC-AG-f02/r02 respectively (**Table S5**). The amplified fragments were cloned via Gibson assembly (using NEBuilder HiFi DNA Assembly Master Mix; New England BioLabs, cat. No. E2621L) into the SbfI/AgeI site of the pLS-SceI vector (Addgene, cat. No. 137725, a gift from Ahituv’s lab) to construct the library. The resulting library was digested with I-SceI (New England BioLabs, cat. No. R0694S) to remove any vector that did not receive an insert. The recombination products were then electroporated into electrocompetent cells (NEB 10-beta, New England BioLabs, cat. No. C3020K) and plated onto Carbenicillin plates. Sanger sequencing of 32 colonies, using n40.dn.F and EGFP.up.R primers (**Table S5**), was then used to confirm the proper assembly of the library. The library was purified using a number of colonies needed to achieve the desired number of Barcodes (n=50) to be associated at each sequence. Barcode-associated fragments were amplified using P7-pLSmp-ass-gfp (100 μM) and P5-pLSmP-ass-i741 primers (**Table S5**), purified using Plasmid plus midi kit (QIAGEN) and tested for its quality via sequencing on a MiSeq (see below section).

### MiSeq

The association between enhancer sequences and the barcodes was ascertained using Illumina MiSeq v2.0 sequencing with pair-ended 150 bp read length. The reads overlap for 30 bp in the middle of the enhancer sequences. Three MiSeq libraries were sequenced to obtain enough number of barcodes covering each tested enhancer sequence with the total number of 168 million reads. The barcodes with 15 bp length were sequenced at the same time with the same read names as the enhancer sequences with the same batch of MiSeq.

### Lentivirus packaging and MOI

The lentiMPRA library was packaged into lentivirus using the plasmid library, psPAX2 (RRID: Addgene_12260) and pMD2.G (RRID: Addgene_12259) and its titration was determined as previously described [22]. In brief, iPSCs were plated at 0.150 million cells/well in 24-well plates and incubated for 24 hours. Serial volume (0, 2, 4, 8, 16, 32 ul) of the lentivirus was added. The infected cells were cultured for 3 days and washed with PBS three times before genomic DNA extraction. Genomic DNA was extracted using the Wizard SV genomic DNA purification kit (Promega). Virus titer and copy number of viral particles per cell were measured by qPCR as previously described [22].

### iPSCs reprogramming and maintenance

Two iPSC lines were used for the lentiMPRA experiment: ACE1815 (two technical replicates, named ACE_1 and ACE_2 in the figures) and 11251. Lines were generated from human skin fibroblasts obtained from a skin biopsy of normal individuals using a viral-free episomal reprogramming method [43, 47] at the Yale Stem Cell Reprogramming Core and passaged for 18 or 20 passages before differentiation. Informed consent was obtained from each donor according to the regulations of the Institutional Review Board and Yale Center for Clinical Investigation at Yale University. The participants agreed to data sharing of genomic unidentified data using controlled data access. All iPSC lines used in this study fulfilled standard reprogramming criteria, including (i) immunocytochemical expression of pluripotency markers (NANOG; SSEA4; TRA1-60); (ii) expression of known hESC/iPSC markers (SOX2, NANOG, LIN-28, GDF3, OCT4, DNMT3B) by semi-quantitative RT-PCR; (iii) downregulation of exogenous reprograming factors. The iPSC lines derived from fibroblasts were cultured on Matrigel (Corning Matrigel Matrix Basement Membrane Growth Factor Reduced)-coated dishes with mTESR1 media (StemCell Technologies) and propagated using Dispase (StemCell Technologies).

### Lentiviral infection

The lentiMPRA library was transduced into iPSCs at the undifferentiated stage. To provide 75%-80% confluency the next day, iPSCs were seeded on Matrigel-coated 10 cm dishes in mTeSR1 media supplemented with 10 uM Y-27632. After 24 hours, the cells were infected with the lentivirus library at an average of Multiplicity Of Infection (MOI) of 4-5 and incubated for 3 days with daily media changes to remove non-integrated virus. The experiment involved three independent replicate cultures, including two lines from different individuals (ACE1815 and 11251), and one technical replicate (ACE1815-1 and ACE1815-2). All cultures were infected at the same time and using the same lentiviral library.

### Forebrain Organoid differentiation

iPSC lines were differentiated into forebrain organoids as described in Jourdon et al 2023 [43]. Briefly, undifferentiated iPSC colonies were treated with 5 μM of the Y27632 compound and dissociated to single cells with Accutase (Millipore, 1:2 dilution in PBS 1X). Four million dissociated cells were seeded in each well of a 6-well plate and cultured in mTeRS1 with 10 μM Y27632 compound on an orbital shaker at a speed of 95 rpm. Forebrain neural induction was triggered by dual SMAD inhibition in mTeSR1 media supplemented with 10μM SB431542, 1μM LDN193189 and 5μM Y-27632 (day1). At day 2, embryoid bodies were cultured in KSR medium (DMEM supplemented with 15% Knockout Serum Replacement, 1% L-Glutamine, 1% NEAA, 1% Pen/Strep and 55 μM of 2β-Mercaptoethanol, 2-ME) with the addition of SB431542, LDN193189, XAV939 and Y-27632. The neural induction with dual SMAD inhibition was maintained until day 7 after which organoids were gradually adapted to NIM medium (DMEM/F12, 1% N2 supplement, 2% B27 without vitamin A, 1% NEAA, 1% Pen/Strep, 0.15% Glucose and 1% Glutamax) through a dilution series of KSR and NIM. Neural progenitors proliferation was induced at day 9 in NIM 75% and KSR 25% supplemented with FGF2 (10ng/ml) and EGF (10ng/ml) and organoids were mantained in proliferative medium until day 16 in 100% NIM. Terminal differentiation was initiated at day 17 (TD0) in terminal differentiation medium (Neurobasal medium supplemented with 1% N2, 2% B27 without vitamin A, 15 mM HEPES, 1% Glutamax, 1% NEAA and 55 μM 2-ME) with the addition of the neutrophic factors BDNF (10 ng/ml) and GDNF (10 ng/ml) until TD30. In the differentiation phase, half of the medium was changed twice a week. Organoids were transferred from wells of a 6-well plate to a 10 cm dish between TD5 and TD10 and the speed of the orbital shaker was decreased to 80 rpm.

### Cell harvesting, library preparation and DNA/RNA extraction and sequencing for barcodes count

The infected cells were harvested at three different time points: in iPSCs after 3 days from the infection before starting organoid differentiation (iPSC stage), and in iPSC-derived forebrain organoids at earlier (TD0) and more mature (TD30) terminal differentiation points. Genomic DNA and total RNA were simultaneously extracted using the AllPrep DNA/RNA Mini Kit (Qiagen, cat. No. 80204) following the manufacturer’s protocol. RNA samples were treated with Turbo DNase (Life Technologies, cat. No. AM1907) to remove any residual DNA contamination. Sequencing libraries were prepared as previously described [22]. Briefly, at least 60 μg total RNA per sample was used for reverse transcription with SuperScript II (Life Technologies, cat. No. 18064-071) using the primer P7-pLSmP-ass16UMI-gfp (**Table S5**) to add a 16-bp UMI and a P7 flowcell sequence downstream of the barcode. PCR steps were performed on the DNA and RNA samples in order to amplify barcodes, adding P5 flowcell sequence and sample index upstream, and P7 flowcell sequence and UMI downsteam to the barcode. Finally, the sequencing libraries were pooled and subjected to paired-end sequencing with UMI, and sample index read.

### Immunostaining

Organoids were randomly selected and fixed in 4% PFA in PBS for 2-4 hours. The organoids were then cryopreserved in 25% sucrose overnight, embedded in O.C.T. (Sakura), and frozen on dry ice before being stored at -80°C. Serial cryosections were obtained with a thickness of 12-16 μm. Immunostaining was performed by incubating the sections in blocking solution (PBS, 10% Donkey Serum, 1% Triton-100) for 1 hour, followed by incubation with primary antibodies (overnight, 4°C) and secondary antibodies (1-2 hours, from Jackson ImmunoResearch or ThermoFisher Scientific). The slides were then mounted with coverslips using VECTASHIELD (Vector Labs) and imaged on a Zeiss microscope equipped with an apotome module and ZEN 3.3 (ZEN pro) software. Three cell lines were used for immunocytochemical analyses, and a minimum of four organoids per line were analyzed. Images were acquired randomly to cover the entire extent of the organoid. Antibody list: FOXG1 (rabbit, 1:200, Takara) and PAX6 (mouse, 1:200, BD Bioscience), EOMES (rabbit, 1:1000, Abcam), FOXP2 (goat, 1:200, Santa Cruz), GAD1 (mouse, 1:200, Chemicon), HuC/D (mouse, 1:200, Invitrogen), SOX1 (goat, 1:100, R&D Systems), CTIP2 (rat, 1:500, Abcam).

### LentiMPRA computational pipeline

#### Pre-processing using MPRAflow and MPRAnalyze

The association between enhancer sequences and barcodes was identified using MPRAflow association package [22] with the following command: nextflow run association.nf --fastq-insert “R1.fastq.gz” --fastq-bc “barcode.fastq.gz” --design “design.fa” --name “MPRAflow” --fastq-insertPE “R2.fastq.gz” -w {workdir} --labels “labels.txt” --cigar 270M.

The count for number of reads for each RNA and DNA barcodes in all 9 samples for all enhancers were calculated using MPRAflow count package [22] with the following command:

nextflow run count.nf --dir “DNA_RNA” --e “experiments.csv” --design “design.fa” --association “MPRAflow_filtered_coords_to_barcodes.pickle” --labels “labels.txt” --umi-length 15 --name “countUMI” --outdir {workdir} –mpranalyze

MPRAnalyze [48] was then performed to normalize the count for RNA barcodes and DNA barcodes in all 9 samples to generate the enhancer activity (RNA/DNA ratio) using the standard pipeline.

#### Identification of negative control distribution using negative references

A mixed Gaussian distribution was applied to calculate the distribution of negative controls using the 135 negative control activity in 9 samples. The curve fit function in Scipy package [49] was applied to estimate the mean and standard deviation of the negative control distribution. The distribution accurately described the negative control data with Kolmogorov–Smirnov test p-value = 0.33 and Anderson-Darling test p-value = 0.12.

#### Candidate regulatory sequences activity quantification and validation

The p-value for the activity of an enhancer to be higher than the negative control distribution was calculated using the enhancer activity (RNA/DNA) and the negative control distribution. The significantly active enhancers were then identified using the p-value of 0.05 after Bonferroni correction using the total number of enhancers tested.

#### Estimating sensitivity of MPRA experiment to define an enhancer as an active one

Using the negative control Gaussian distribution, we defined a cutoff for the RNA/DNA ratio that corresponds to a significant p-value. We then calculated the area under the positive control Gaussian and above the cutoff. The sensitivity was then estimated as a ratio of the area and the total area under the positive control Gaussian distribution.

### Clustering of enhancer activity for candidate regulatory sequences

Enhancer activity (RNA/DNA) of candidate regulatory sequences and positive reference sequences was calculated by MPRAnalyze. Subsequently, Seaborn 0.11.0 [50] clustermap was employed for clustering the enhancer activity values. Ward’s minimum variance method was utilized for clustering. The resulting heatmap displayed the clustered enhancer activity values. The color used for enhancer activity larger than 4 was the same as that used for the value of 4.

### Transcription factor binding site identification and expression of the transcription factors

FIMO [33] was employed to identify TFBSs from all 7,289 sequences, including candidate regulatory sequences, positive reference sequences, and negative reference sequences. The transcriptional factor binding site annotation was downloaded from JASPAR2022 [51] with 1,252 Homo Sapiens annotations. The command for running FIMO [33] is as follows:

fimo --o {out_dir} JASPAR2022_tfbs input.fa.

TFBSs identified by FIMO with FDR <= 0.05 were taken for further analyses. We quantified the number of TFBSs in each sequence and compared the distribution between validated and non-validated candidate regulatory sequences using Kolmogorov–Smirnov test. We further calculated the expression (RPKM) of the TFs which bind to these TFBSs in 97 TD0/TD30 organoid samples. The expression of TFs in validated and non-validated candidate regulatory sequences was compared using Kolmogorov–Smirnov test.

### Genes regulated by candidate regulatory sequences and expression

Genes regulated by candidate enhancers were identified by Amiri et al. [23] taking confident_set1, confident_set2, and proximity genes. The expression (RPKM) of the genes was calculated from 97 TD0/TD30 organoid samples.

### Intersection with external datasets

Intersection with external datasets were performed using BedTools [52] and the resulting data was plotted using the ‘UpSetR’ package in R [53].

### Single cell ATAC-seq analysis

For each 10X scATACseq sample, fastq files were first processed by cellranger-atac v2.0.0 with default parameters and 10X prebuilt reference arc-GRCh38-2020-A-2.0.0. The resulted cell-by-peak count matrix was first processed and filtered by Signac following online vignette (https://stuartlab.org/signac/articles/pbmc_vignette.html). Cell types were annotated using our annotated scRNAseq dataset and the label transfer method implemented in Seurat following online vignette (https://satijalab.org/seurat/articles/atacseq_integration_vignette.html). scATACseq peaks were then called using all reads in a sample or subsets of reads from each annotated cell type by running Signac function CallPeaks with default parameters.

## Acknowledgements

We thank the members of the Vaccarino lab for extensive discussions, technical help and contributions to methods. We thank Guilin Wang and Christopher Castaldi, and the Yale Center for Genome Analysis for library preparation, deep sequencing, and Cell Ranger analysis. We thank Caihong Qiu and Jason Thomson at the Yale Stem Cell Center for the generation of the iPSC lines. Part of the illustrations were created using BioRender. Com.

## Data sharing

The source MPRA data described in this manuscript are available via the PsychENCODE Knowledge Portal (https://psychencode.synapse.org/). The PsychENCODE Knowledge Portal is a platform for accessing data, analyses, and tools generated through grants funded by the National Institute of Mental Health (NIMH) PsychENCODE Consortium. Data is available for general research use according to the following requirements for data access and data attribution: (https://psychencode.synapse.org/DataAccess). For access to content described in this manuscript see: https://doi.org/10.7303/syn51072171.1. The source bulk RNA-seq data are available at the NIMH Data Archive (NDA) under collection #C2424, at url: https://nda.nih.gov/edit_collection.html?id=2424.

## Supporting information

**S1 Fig. Minimal enhancer region selection**.

**S2 Fig. Immunocytochemical characterization of forebrain organoids**.

**S3 Fig. Statistics of MiSeq barcodes for tested enhancers**.

**S4 Fig. Gaussian mixture model with positive and negative controls**.

**S5 Fig. RNA/DNA ratios in all tested enhancers at iPSC, TD0 and TD30 comparing between different samples**.

**S6 Fig. Intersection with external datasets**

**S7 Fig. MPRA-active enhancers positively or negatively correlate with expression of the corresponding linked genes**.

**S8 Fig. MPRA-active enhancers correlate with differences in expression of linked genes across timepoints**.

**S9 Fig. MPRA-active enhancers positively or negatively correlate with expression of the corresponding linked genes**.

**S1 Table. MPRA-tested enhancers activity across samples**.

**S2 Table. Intersection of MPRA-tested enhancers with external datasets**.

**S3 Table. Enhancers activity, TFs and linked-genes expression**.

**S4 Table. Intersection of MPRA-tested enhancers with scATAC-seq**

**S5 Table. Primers sequences for library amplification, virus titration and sequencing**

